# Adaptation of *Mycobacterium tuberculosis* to biofilm growth is genetically linked to drug tolerance

**DOI:** 10.1101/663369

**Authors:** Jacob P. Richards, Wenlong Cai, Nicholas A. Zill, Wenjun Zhang, Anil K. Ojha

## Abstract

*Mycobacterium tuberculosis* (Mtb) spontaneously grows at the air-medium interface forming pellicle biofilms, which harbor more drug tolerant persisters than planktonic cultures. The underlying basis for increased persisters in Mtb biofilms is unknown. Using a Tn-seq approach, we show here that multiple genes that are necessary for fitness of Mtb cells within biofilms, but not in planktonic cultures, are also important for their tolerance to a diverse set of stressors and antibiotics. Thus, development of Mtb biofilms appears to be associated with population enrichment, in which endogenous stresses presumably generated by challenging growth conditions within biofilm architecture select for cells that maintain tolerance to exogenous stresses including antibiotic exposure. We further observed that the intrinsic drug tolerance of constituent cells of biofilms determines the frequency of persisters: morphologically indistinguishable monoculture biofilms of a Δ*pstC2A1* mutant hypersensitive to rifampicin harbor ∼20-fold fewer persisters than wild-type. These findings together allow us to propose that the selection of elite cells during biofilm development significantly contributes to the persister frequency. Furthermore, probing the possibility that the population enrichment is an outcome of unique environment within biofilms, we demonstrate biofilm-specific induction in the synthesis of isonitrile lipopeptides (INLP). Mutation analysis indicates that INLP is necessary for the architecture development of Mtb biofilms. In summary, the study offers an insight into persistence of Mtb biofilms under antibiotic exposure, while identifying INLP as a biomarker for further investigation of this phenomenon.

**SIGNIFICANCE:** The tuberculosis (TB) pathogen *Mycobacterium tuberculosis* (Mtb) is one of the deadliest bacterial pathogens known to mankind, and TB treatment is inefficient. A lengthy chemotherapy for TB is attributed to a small subpopulation of Mtb bacilli exhibiting phenotypic tolerance to antibiotics. Drugs targeting these persisters are expected to shorten TB chemotherapy, but their development is dependent on *in vitro* growth models that reproducibly generate high frequency of persisters. Biofilms of Mtb are a suitable model for understanding the origin of persisters. Here, we provide an explanation for the elevated persister frequency in Mtb biofilms. We also identify isonitrile lipopetides as a biomarker of Mtb biofilms. These findings will facilitate further advancements of our efforts to identify and target Mtb persisters.

## INTRODUCTION

Treatment of tuberculosis (TB), caused by *Mycobacterium tuberculosis* (Mtb), entails a multi-drug regimen administered for at least 6 months. A lengthy treatment is presumably necessitated by the persistence of a small subpopulation of bacilli, which exhibit phenotypic tolerance to antibiotics (1, 2). Although the mechanism underlying the development of Mtb persisters during infection is unclear, *in vitro* studies suggest that these bacilli probably develop through both stochastic and induced mechanisms(3–6).

The persistence of microbes against antibiotics has been closely linked to their ability to grow as sessile, three-dimensionally organized, matrix encapsulated, multicellular communities called biofilms (7–11). Biofilm-related antibiotic tolerance is rendered particularly relevant by the fact that a majority of chronic microbial infections in humans occur as biofilms (12). Growth and development of biofilms is a multi-stage process that requires dedicated genetic programs expressed in a spatiotemporal order(13–15). Initial stages involving substratum attachment of planktonic cells and aggregated growth are followed by a maturation stage accompanied by synthesis of an extracellular matrix (13, 14). Physical contacts and chemical communications among cells in developing biofilms, as well as physiological adaptation to self-generated gradients of nutrients and oxygen stratify the architecture resulting in phenotypically heterogeneous cells (15–18). Moreover, molecular mechanisms of stress and antibiotic tolerance in biofilm cells are considered to be linked to those involved in physiological differentiation during architecture development (16, 19).

Environmental and pathogenic mycobacteria, like *Mycobacterium avium*, *Mycobacterium abscessus*, *Mycobacterium smegmatis* and *Mycobacterium tuberculosis* spontaneously form biofilms under detergent-free *in vitro* growth conditions (20–26). Biofilms of *M. avium*, *Mycobacterium abscessus*, *Mycobacterium fortuitum* and *Mycobacterium ulcerans* have also been reported in environmental specimens and in host tissues (27–30). Mutants of mycobacteria that exhibit defective biofilm development, without any defect in planktonic growth, demonstrate the requirement for specialized genes for biofilm development. Moreover, the frequency of drug tolerant persisters in Mtb biofilms are higher than in planktonic cultures, and their occurrence is tightly linked to the development of a mature three-dimensional architecture (22, 31, 32), similar to that observed in other bacterial species. A previous evidence of the involvement of GlnR in nitrogen assimilation as well as peroxide resistance during biofilm development of *M. smegmatis* offers interesting molecular insights into the biofilm-associated stress tolerance (33), although such ideas have been largely unexplored in Mtb.

Although biofilms of Mtb during infection remain to defined, *in vitro* biofilms represent a suitable model for understanding the origin of persisters. In this study, we report that genes that confer fitness of Mtb within biofilms are also involved in maintaining tolerance to antibiotics, suggesting that the growth environment in biofilms selects for cells that maintain intrinsic tolerance to exogenous stress including antibiotics. We further show that intrinsic drug tolerance of the constituent cells within biofilms is a determinant of the persister frequency. Lastly, we discover that induced synthesis of isonitrile lipopetides (INLP) is specific to Mtb growth in biofilms, thus demonstrating the unique environment generated in biofilms. These findings begin to explain the basis for biofilm-associated antibiotic tolerance in Mtb.

## RESULTS

### Tn-seq of Mtb biofilms

To comprehensively determine the genes that confer a fitness advantage to Mtb in biofilms, we employed a previously developed high-throughput genetic screen, Tn-seq (34). We cultured a transposon insertion mutant library of Mtb planktonically and in pellicle biofilms (Fig. 1a). Although Sauton’s medium is more suitable for formation of wild-type Mtb biofilms (22), we chose nutrient-rich 7H9OADC medium for Tn-Seq screening to minimize a potential growth bias introduced due to selective nutrients of Sauton’s medium. Sauton’s medium however was used for all subsequent monoculture studies of genes identified from the Tn-Seq screen. We also analyzed colonies of the Tn-library by Tn-seq (Fig. 1A). While colonies and pellicles represent different growth models *in vitro*, the two have overlapping gene expression patterns in *M. smegmatis*(14). Inclusion of both models in the Tn-seq screen allowed similarities and differences in genetic requirements for Mtb fitness to be determined in these two models of aggregated growth. We recognize that Tn-seq has limitations for screening the colony model of growth, because the procedure involves post-exposure outgrowth of individual mutants on agar plates (34). Mutants with an inability to form clonally pure colonies are expected to be eliminated in this screen. Thus, the screen is limited to only those mutants that can form normal colonies in monocultures, but display fitness loss under direct competition with wild-type. Furthermore, Tn-seq can also potentially fail to report mutants that are trans-complemented by wild-type in mixed biofilms. Nevertheless, Tn-seq remains a powerful genetic screening approach for fastidious organisms like Mtb.

**Figure 1:**
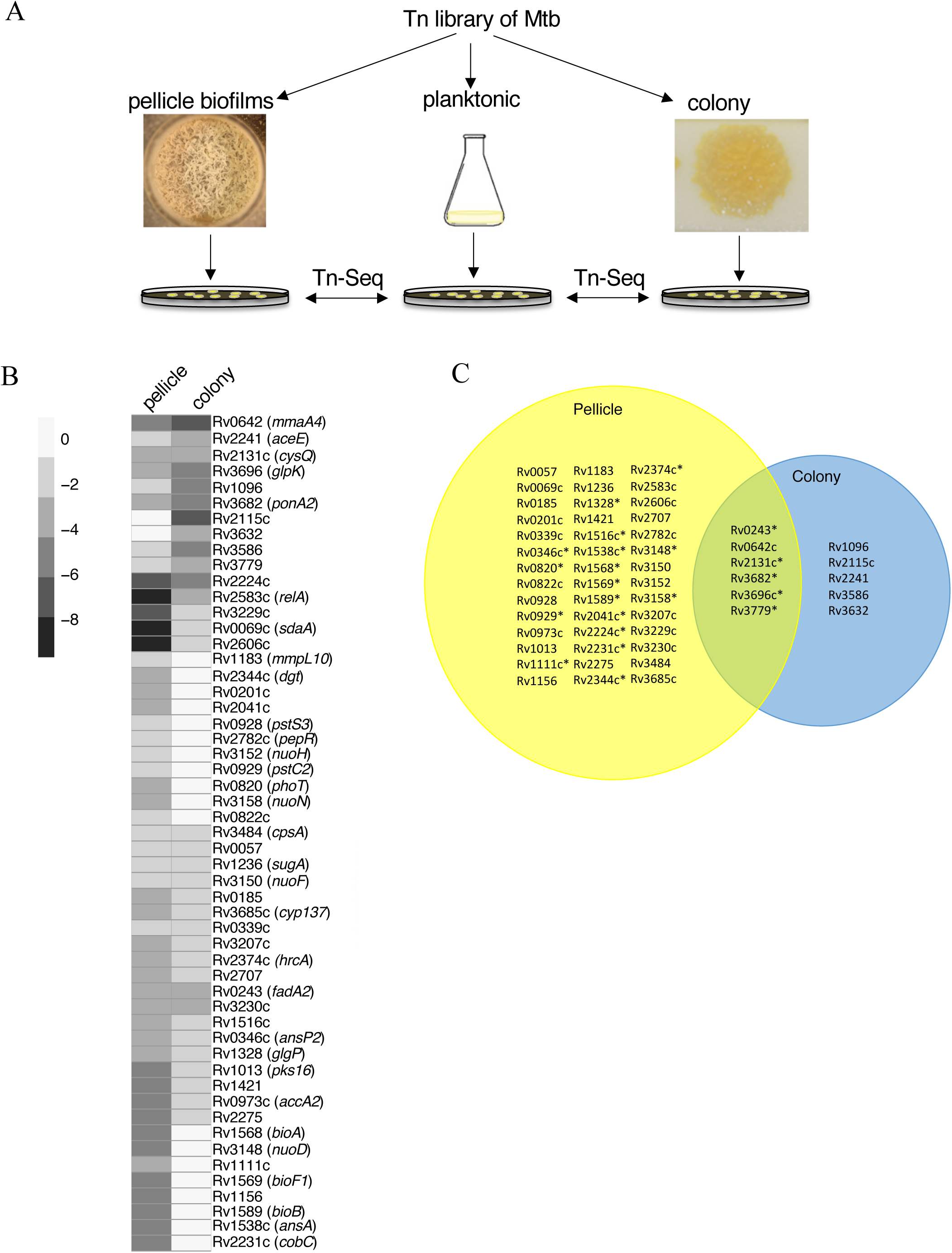
Genes necessary for Mtb fitness within biofilms. **A.** A summary of Tn-seq based approach to identify genes required for fitness of Mtb (mc^2^7000) in colonies or pellicle biofilms, relative to planktonic cultures. **B-C.** Heat map (B) and Venn diagram (C) of 53 genes, mutations in which cause significant underrepresentation of clones either in pellicle biofilms or colonies. While the heat map depicts fold-change in the reads, the Venn diagram depicts statistically significant read counts in each of the two growth models. The scale bar for the heat map represents log_2_(fold-change). Fold changes of six genes (Rv# 0057, 0339, 1236, 2131c, 3150 and 3484), depicted by the heat map as similar in both colonies and pellicles (B, are statistically significant only in the pellicle model (C). The metadata summary of Tn-seq is provided in Table S1, whereas complete dataset obtained from TRANSIT analysis is in Table S2. Asterisks indicate genes that were followed-up in this study.

Using massively parallel sequencing, we compared the frequency of transposon junction site per gene in colonies and biofilms relative to planktonic cultures. Reads corresponding to junction sequences of transposon from each growth condition were obtained through sequential processing of filtering and trimming using the TRANSIT pipeline (35). Of the total 74,605 potential transposon insertion sites (TA), the mapped TA sites in samples ranged from 29.5% to 55.7% coverage (Table S1). However, no correlation was observed between growth conditions and density of mapped reads (Table S1). Only one sample had a library density below the recommended threshold of 35% coverage (35). One of the replicates in the colony model had the least (29.5 %) library density, and also returned significantly less reads than the other sequencing libraries (Table S1). Removing this replicate from TRANSIT did not affect the comparison of Tn hits between colonies and planktonic cultures; Pearson correlation coefficients between the sequencing replicates for each growth condition indicated statistically significant consistency (Figure S1).

We identified 53 candidate genes that were significantly (q-value < 0.05) underrepresented in either pellicles or colonies, or both, relative to planktonic cultures (Fig. 1B-C and Table S2). Of these, 48 mutants were under-represented exclusively in pellicle biofilms, while 11 were under-represented in colony biofilms. The differences in fitness patterns of the mutants between pellicles and colonies suggest a distinct genetic requirement for growth under each condition. The list of 53 genes included *mmaA4*, *pks16* and *relA* (Fig. 1B-C), which were previously implicated in formation of pellicle biofilms (22, 31, 36). Consistent with the altered morphology of Δ*mmaA4* colonies, and its deficiency in formation of pellicle biofilms (31), Tn mutants of *mmaA4* were underrepresented in both colonies and pellicles (Fig. 1B-C).

### Monocultures of mutants distinguish absolute from relative fitness deficiency

Excluding the previously reported *mma4*, *relA* and *pks16* genes, we selected 22 candidates (indicated in Fig. 1C) for investigation of their role in biofilm formation. The selection was based on known significance of these genes in growth and/or pathogenesis of Mtb (34). Isogenic deletions in these genes were constructed and their phenotypes in monoculture pellicle biofilms were compared with their planktonic growth. The comparison revealed three categories of mutants. In the first and the largest category, 16 mutants were indistinguishable from the wild-type strain in their ability to form biofilms, suggesting that mutations in these genes exhibit fitness deficiency only when they are in direct competition with wild-type in biofilms. In the second category, two mutants (Δ*ansA* and Δ*bioA*) were severely growth retarded in both planktonic and biofilm cultures (Fig. 2A-C). This suggests that growth in Sauton’s medium, which contains L-asparagine as the primary nitrogen source and lacks supplemental biotin, requires the activity of L-asparaginase (*ansA*) and biotin synthase (*bioA*) in cells. The activities of *ansA* and *bioA* may have greater consequence in biofilm formation than in planktonic growth because mutations in these genes cause a biofilm-specific fitness disadvantage in the Tn-Seq screen, which was performed in biotin- and nitrogen-rich 7H9OADC medium. The last category of four mutants formed either morphologically altered (ΔRv2224c and Δ*ponA2*), delayed (Δ*phoT*), or severely deficient (Δ*dgt*) pellicle biofilms, but remained growth competent in planktonic cultures (Fig. 2A-B). The Δ*phoT* mutant only formed normal biofilms upon extended incubation (Figure S2). The Δ*dgt* phenotype was particularly striking because the mutant accumulated biomass at the bottom of the container (Fig. 2A). Plasmid-borne expression of *dgt* under a constitutive *hsp60* promoter partially complemented Δ*dgt* (Fig. 2A), suggesting that additional chromosomal elements are perhaps necessary. Slow development of Δ*phoT* biofilms was also rescued by plasmid-borne expression of *phoT* by the *hsp60* promoter (Fig. 2A). All four mutants exhibiting impaired pellicle development were able to grow as colonies, but with altered morphology (Figure S3).

**Figure 2:**
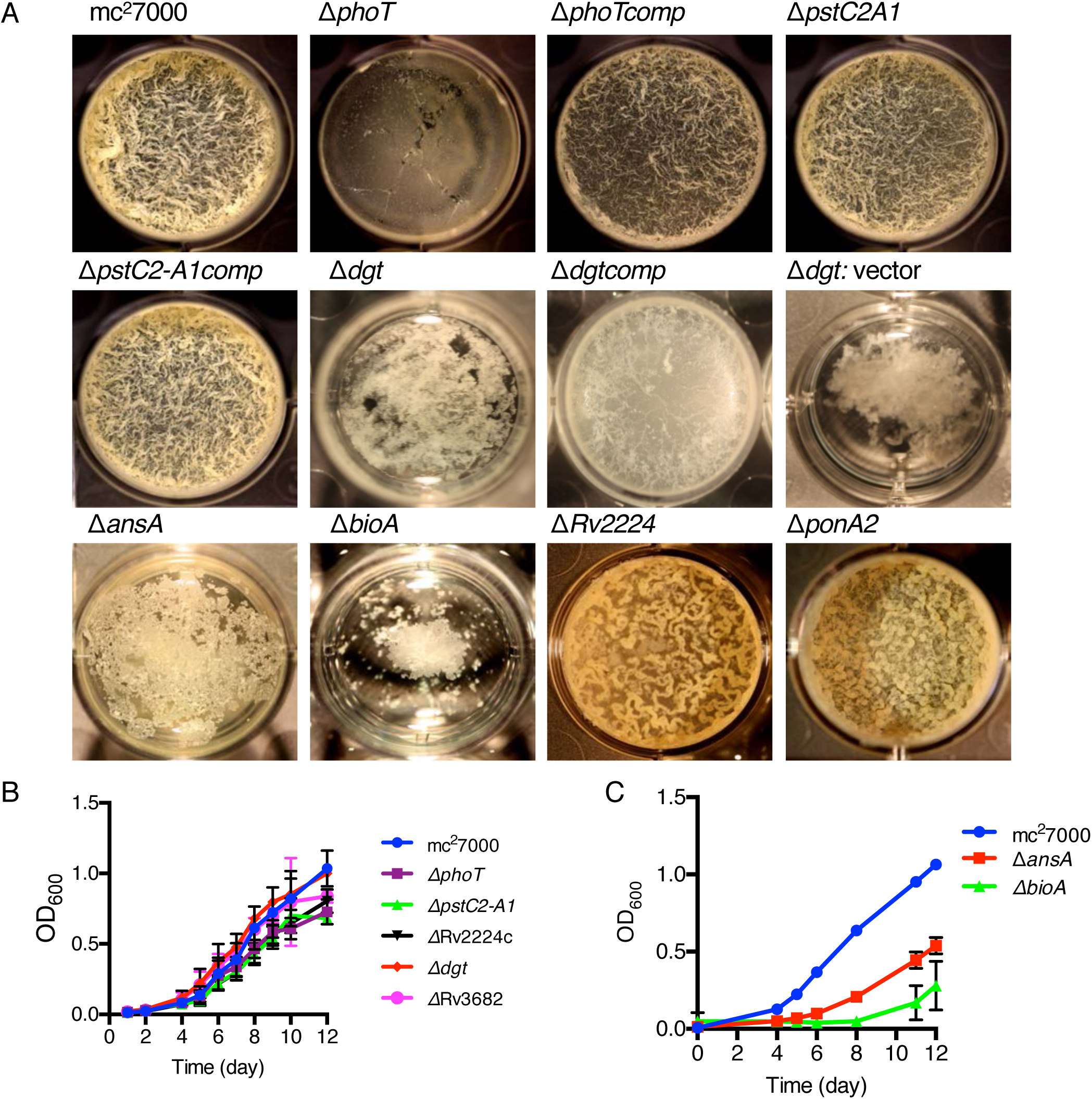
Genes required for development of *M. tuberculosis* in pellicle biofilms. **A.** A top down view of pellicles of mc^2^7000 (wild-type) and a subset of isogenic mutants of genes identified from the Tn-seq screen (see Fig.1C). Complemented strains of Δ*phoT (*Δ*phoTcomp)*, Δ*pstC2A1 (*Δ*pstC2A1comp)*, Δ*dgt (*Δ*dgtcomp)* carrying a plasmid expressing the corresponding genes under the control of either its own promoter (for *pstC2A1)* or the *hsp60* promoter are also shown. The strain Δ*dgt* carrying the empty vector pMH94 was control for Δ*dgtcomp*. Pellicles were grown at the air-medium interface in 12-well tissue culture plates in Sauton’s medium for 5 weeks at 37°C. While the biomass of Δ*dgt* remained at the bottom of the well, Δ*phoT was* delayed and formed normal pellicles upon further incubation (see Figure S2). **B-C.** Planktonic growth of mc^2^7000 and the mutant strains described in figure 2A. All strains were cultured in detergent-free Sauton’s medium. Cultures were shaken once daily, and their optical densities were measured after dispersion of 1mL aliquots in Tween-80. Data represent mean ± SD (n=3).

The low abundance of *pstC2A1* and *phoT* mutants in biofilms of the Tn-library reveal the importance of phosphorus (Pi) homeostasis in fitness of Mtb cells within biofilm microenvironments. Given that Rv0928-30 encoded PstS3, PstC2, PstA1, and Rv0933 encoded PstB constitute the active Pi-transporter complex (37, 38), a rather mild impairment of Δ*pstC2A1* in monoculture biofilms relative to Δ*phoT* was unexpected. An RNA-seq analysis of the two mutants in comparison with wild-type offers a possible explanation. The transcriptomic profiles of Δ*pstC2A1* and Δ*phoT* exhibited extensive similarity in the patterns of gene expression relative to wild-type (Figure S4 and Table S3), consistent with the idea that both the loci indeed function in Pi homeostasis. RegX3, a transcription regulator that responds to Pi depletion through its cognate sensor SenX3 (39), and the downstream target operon *pstS3C2A1* were upregulated in both the mutants (Table S3). Expression levels of PhoT remained constitutively low in all the strains, but transcripts corresponding to its putative paralogue encoded by Rv0933 were constitutively higher than PstS3C2A1(Table S3). Therefore, the upregulation of *pstS3C2A1* operon and high constitutive levels of PstB (Rv0933) in Δ*phoT* would increase the levels of functional PstS3C2A1B complex, thereby increasing the abundance of active Pi transporters causing dysregulation of Pi homeostasis. This scenario is inconceivable in the Δ*pstC2A1* mutant, which may perhaps less severe dysregulation of Pi homeostasis than Δ*phoT*.

### Relationship between fitness in biofilms and persistence against antibiotics

Multiple genes of diverse functional categories (like *mma4*, Rv2224c, *ponA2*, and *pstC2A1*) identified in our Tn-seq screen have also been implicated in maintaining intrinsic tolerance to a panel of stressors and antibiotics, including but not limited to acidic pH, lysozyme, SDS, isoniazid and rifampicin (31, 40, 41). A reasonable interpretation of this observation is that the restrictive conditions formed in biofilms may favor the growth of cells that are tolerant to both endogenous and exogenous stresses. Thus, more cells with intrinsic antibiotic tolerance are expected to occur in biofilm population than in planktonic cultures. Given that intrinsic defense mechanisms against antibiotic-induced damages are closely linked to increased persister frequency in a bacterial population (42), we hypothesize that enrichment of drug tolerant cells in Mtb biofilms could also impact the persister frequency (Fig. 3A). A corollary to this hypothesis is that monoculture biofilms of mutant cells that are intrinsically sensitive to antibiotics harbor fewer persisters than wild-type. To test this, we exploited the Δ*pstC2A1* strain, which exhibits hypersensitivity to rifampicin (RIF) (40). Moreover, the mutant forms biofilms indistinguishable from wild-type, thereby allowing a direct assessment of the effect of intrinsic property of the cells without any compounding effect from an altered architecture, which is generally observed with other antibiotic sensitive mutants identified in this study (e.g. *ponA2*, *mma4*). Matured biofilms of the Δ*pstC2A1* mutant and wild-type strains were exposed to 50 μg/mL of RIF, which is 100-times the minimal inhibitory concentration, for 7-days, and the persister frequency was scored as described previously (22). Relative to wild-type, the mutant biofilms contained significantly (∼20-fold) fewer RIF tolerant persisters. The persister frequency was substantially restored upon complementation (Fig. 3B-C). A mutation in *pstA1* also produces fewer persisters in planktonic cells (40), suggesting that regardless of the growth condition, intrinsic drug sensitivity of the representative population is a key determinant of the persister frequency. Therefore, more drug tolerant cells in biofilms than in planktonic culture increase the frequency of persisters.

**Figure 3:**
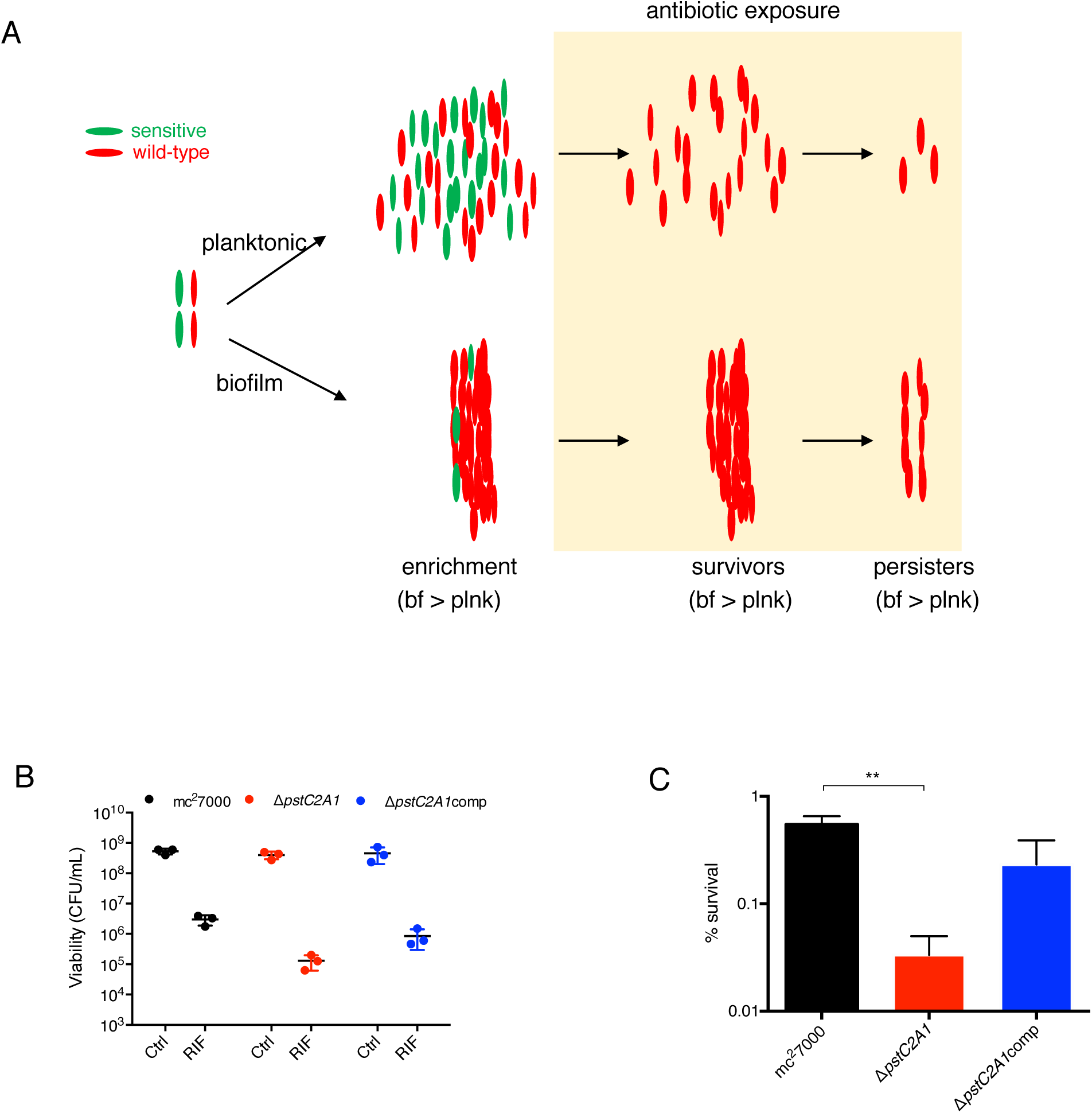
Intrinsic antibiotic tolerance of constituent cells determines persister frequency in Mtb biofilms. **A.** A schematic representation of biofilm-associated drug tolerance in Mtb. Enrichment of wild-type (red) over sensitive (green) cells in biofilms increase the frequency of persisters. Thus, a monoculture biofilms of a hypersensitive mutant is expected to harbor fewer persisters than wild-type. **B-C.** Frequency of rifampicin tolerant persisters in biofilms of mc^2^7000, a RIF sensitive Δ*pstC2A1* and Δ*pstC2A1comp.* Five week pellicles were exposed to 50 *µ*g/mL RIF for one week, then homogenized with Tween-80 and dilutions were plated on 7H11 agar plates to enumerate viable cells. Cells exposed to DMSO (solvent for rifampicin) were used as untreated control (Ctrl). Data represent three biologically independent experiments. Average percentage (%) survival calculated from panel B plot are shown in panel c (n= 3; p < 0.01; unpaired t-test).

### Unique growth environment in biofilms is marked by INLPS induction

We hypothesize that the enrichment of stress/antibiotic tolerant cells in biofilms likely originates from their complex microenvironment that emerge from the uniquely stratified architecture. To demonstrate a distinctive growth environment of Mtb biofilms, we sought to identify a gene-expression pattern specific to Mtb growth in biofilms. We compared transcriptomic analysis of planktonic and biofilm cultures (Table S4). Co-induction in biofilms of a 6-gene operon (Rv0096-0101), which is involved in the synthesis of a putative isonitrile lipopeptide (INLP) and thus called INLP synthase (INLPS) complex (Fig. 4A) (43), was immediately apparent (Table S4). We confirmed the induced expression of Rv0096 in 4-week biofilms by RT-PCR, which was not evident during an earlier (2-week) stage (Fig. 4B). Expression of a Dendra2 reporter gene fused to the *inlps* promoter (P*_inlps_*) was only detected at the 4-week stage. By contrast, the reporter expression was not observed at an earlier (2-week) stage of biofilm development, or in planktonic culture, or in colonies grown on an agar surface (Fig. 4C and Figure S5). Reporter expression was further confirmed in 4-week biofilms of the virulent strain of Mtb (Erdman) (Figure S5). Thus, INLPS induction appears to occur specifically during later developmental stages of pellicle biofilms, but not in any other forms of growth. This pellicle-specific INLPS induction further highlights the distinction between pellicle biofilms and colony growth as suggested by the Tn-seq results. INLPS is negatively regulated by RegX3 (44), suggesting that the retarded biofilms of the Δ*phoT* mutant could also be contributed by a suboptimal induction of the cluster. While Rv0096 induction was not apparent in 4-week biofilms of Δ*phoT*, the expression was fully restored upon further incubation (Fig. 4D). This implies that factors other than RegX3, possibly responding to signals generated during maturation of biofilms, likely regulate INLPS. The correlation between INLPS induction and timing of biofilm maturation further suggested an important role of INLP during the maturation stages. A severe defect in biofilm formation of Δ*inlps* mutant was confirmed (Fig. 4E), without any growth defect in planktonic culture (Figure S6). The mutant phenotype was partially restored upon complementation with a cosmid bearing *inlps* and its upstream promoter region (Fig. 4E). The Δ*inlps* mutant formed relatively smooth and shiny colonies, in contrast to dry and rugose textured wild type colonies (Figure S6), indicating that the basal level of INLP expression also has important functional significance.

**Figure 4:**
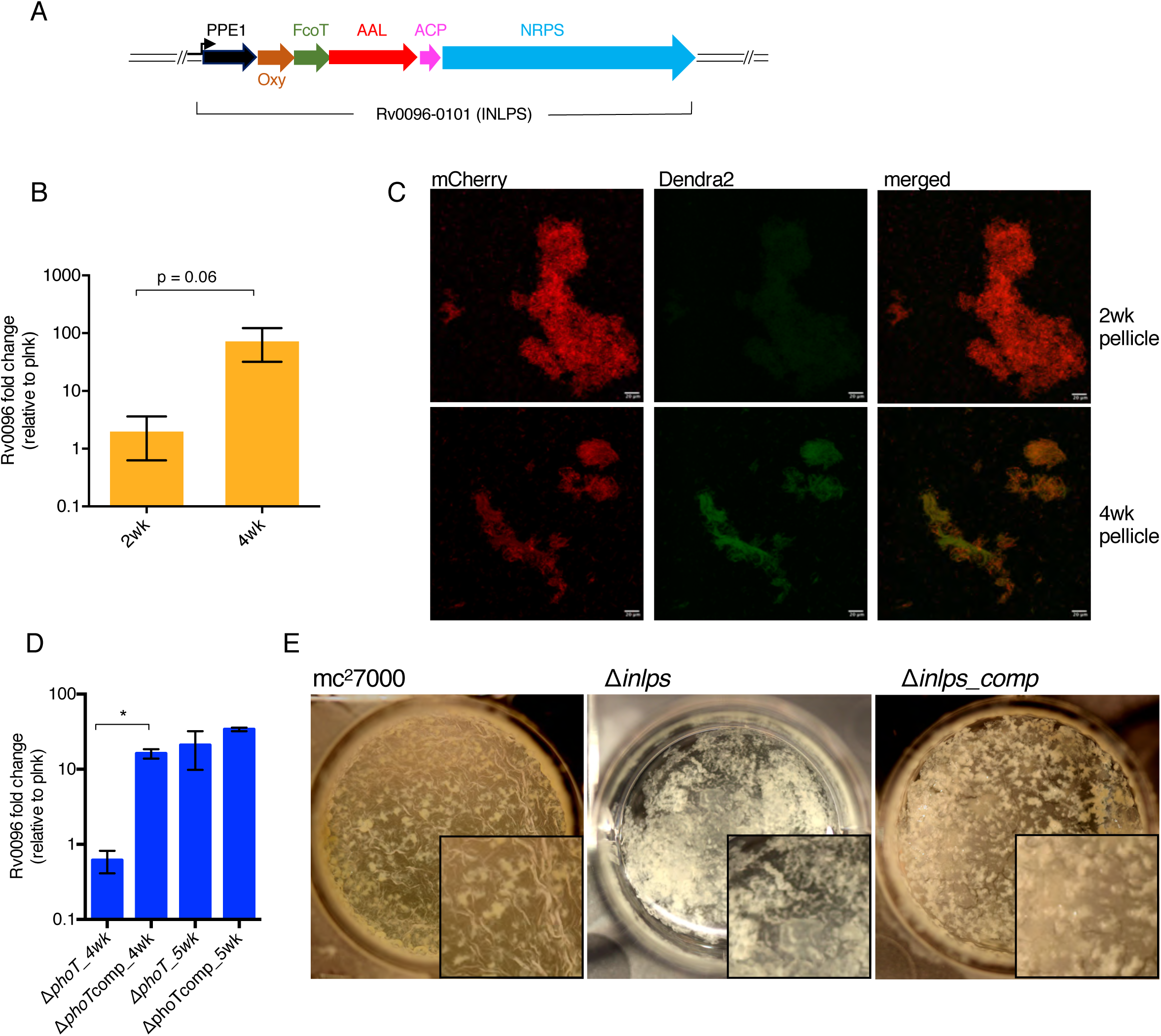
Induced expression of INLPS is required for biofilm development. **A.** Schematic representation of *inlps* gene cluster comprising of six ORFs organized in an operon. Except for Rv0096, which has the conserved Pro-Pro-Glu (PPE) motif, likely function of other genes have been assigned based on previous studies^44^. Oxy: oxidase, FcoT: thioesterase, AAL: acyl ACP ligase, ACP: Acyl Carrier Protein and NRPS: Non-ribosomal peptide synthase. The arrow indicates the putative promoter. **B.** RT-PCR based expression analysis of Rv0096 in mc^2^7000 biofilms at 2-week and 4-week stages, relative to planktonic (plnk) culture. **C.** Microscopic analysis of Dendra2 expression from the promoter of *inlps* in 2- and 4-week stages of mc^2^7000 biofilms. Cells constitutively expressing mCherry from the *hsp60* promoter were smeared in 80% glycerol on a glass slide and images were captured with 20X objective on a confocal microscope, analyzed by FIJI software, and shown as maximum projection. Scale bar corresponds to 20*µ* **D.** An RT-PCR analysis of Rv0096 expression in the *ΔphoT* mutant and a complemented (*ΔphoT*comp) strains at 4- and 5-week stages of pellicle biofilm development. SigA transcripts were used as endogenous control for normalization. Data in panels B and D represent average of at least two biologically independent experiments. * p < 0.05 (paired t-test). **E.** A top-down view of 4-week pellicle biofilms of mc^2^7000, its isogenic Δ*inlps* mutant, and a complemented strain with cosmid-borne *inlps*. While the parent wild-type and the complemented strain had visible texture development at the air-medium interface, the biomass of the mutant was predominantly at the bottom of the well.

### Chemical evidence of INLP accumulation in biofilms

We next sought to determine if expression of *inlps* correlates with new metabolite accumulation in Mtb biofilms. INLP was originally identified upon overexpression of Rv0097-0101 and its homologues in *E. coli* (43), but the structure of the INLP native to Mtb is unknown. We therefore took an unbiased approach and compared lipid profiles of crude organic extract from planktonic and biofilm cultures of wild-type Mtb. The biofilm culture of Δ*inlps* was also included for comparison. The comparative profiling by Liquid Chromatography-High Resolution Mass Spectrometry (LC-HRMS) analysis detected an increased abundance of more than two hundred small molecules in wild-type biofilm extracts relative to other extracts (data not shown), complicating the identification of metabolic products of the *inlps* gene cluster. Further careful analysis of these metabolites idneified four compounds with identical masses (m/z: 717.5612, proposed molecular formulas C_40_H_73_N_6_O_5_^+^), which are likely synthesized by the *inlps* cluster (Fig. 5; **1-4,** and Figure S7). Our analysis was based on three criteria. First, these four compounds were detected exclusively in wild-type biofilm extracts, and were not detected in either planktonic wild-type or in Δ*inlps* cultures (Fig. 5A). Identical fragmentation patterns by LC-MS/MS analysis of these metabolites were observed (Figure S7) indicating that these compounds are isomers. Second, click reactions with tetrazine confirmed the presence of unique isonitrile functionality in the compounds (45) (Fig. 5B-C). Tetrazine treatment of partially purified **1** and **2** caused disapperance of their mass signals with concomitant appearance of the predicted reaction product. Third, weak but noticable incoporation of ^13^C-glycine was observed in these four metabolites (Figure S8). Glycine is a direct precursor utilized by INLPS (43, 45). Therefore its incorporation in 1-4 is in agreement with the characterized biosynthetic mechanism for INLP synthesis. We note that the structure of these molecules remains to be determined, largely due to the difficulty in purification of these molecules in sufficient quantity necessary for NMR analysis. Nonetheless, our data provide substantial evidence that compounds 1-4 are isonitrile lipopeptides, and their accumulation in biofilms is a direct consuequence of induced expression of *inlps*.

**Figure 5:**
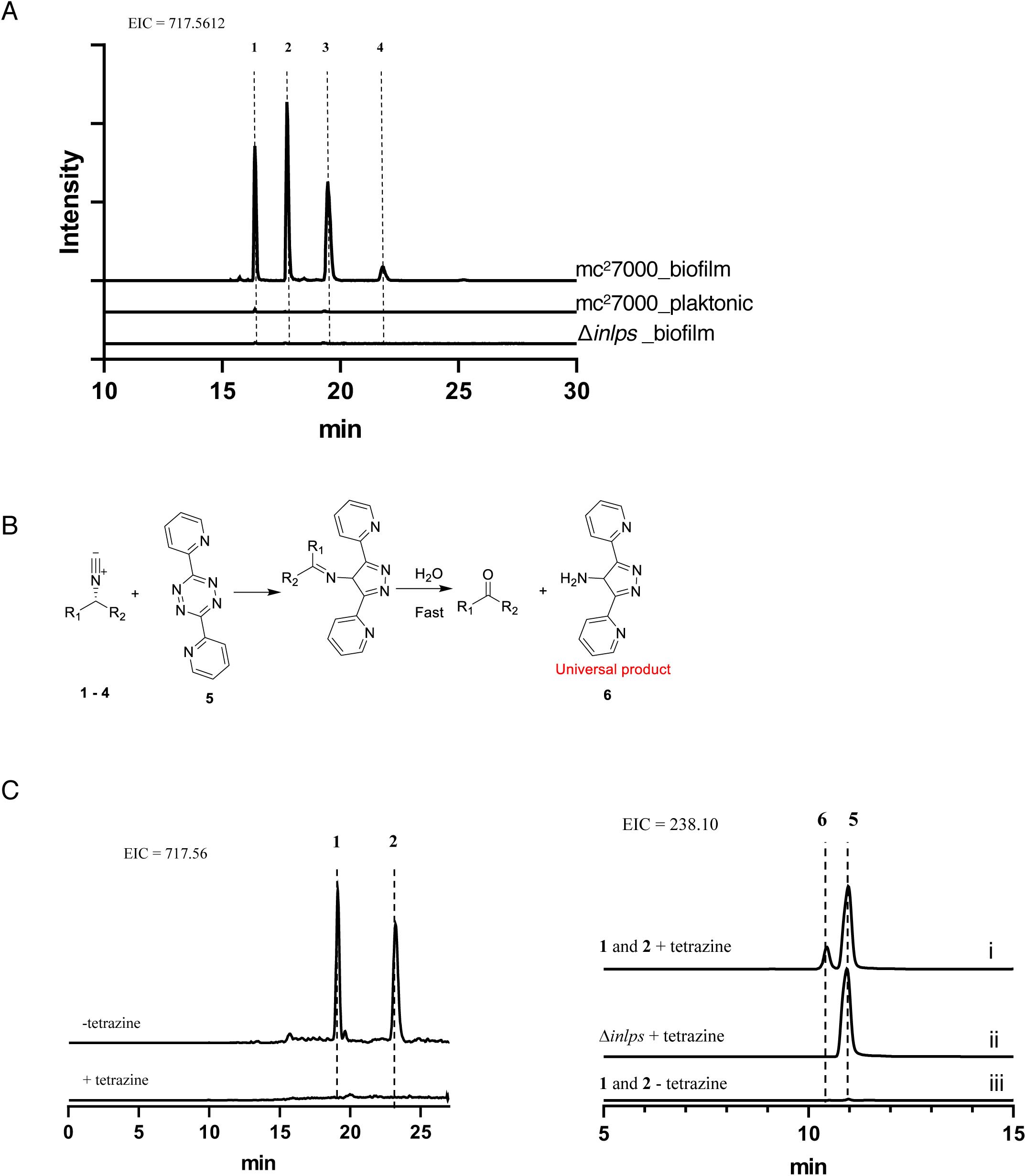
Accumulation of INLP in biofilms of Mtb. **A.** Extracted ion chromatograms (EICs) show presence of four isomers of INLP (**1**-**4;** *m/z* = 717.5612) exclusively in the wild-type biofilm extract, but neither in planktonic wild-type nor in Δ*inlps* cultures. A 10-ppm mass error tolerance was used for each trace. **B**. Scheme of click reaction of INLP with tetrazine showing [4+1] cycloaddition between **1-4** and tetrazine. The product rapidly hydrolyzes in a trace amount of water to generate a primary amine **6**. **C** Extracted ion chromatograms show disappearance of partially purified **1** and **2** after reaction with **5** and concomitant appearance of the predicted reaction product **6.** Control reactions without **5,** and **5**-treated Δ*inlps* mutant extract are also shown.

### Signal inducing INLP originates in biofilm architecture

We next sought to determine the origin of the signal that regulates the *inlps* gene cluster. No changes were detected in RegX3-Senx3 transcripts during biofilm development (Table S3), thereby ruling out the possibility that *inlps* upregulation is signaled by changes in Pi homeostasis. Moreover, the level of *inlps* induction in a *regX3* mutant (44) is about an order of magnitude less than those observed in biofilms (Table S5). We thus hypothesized that the signal regulating *inlps* induction is dependent on the specific nature of growth environment in mature pellicle biofilms, which are not created by other growth conditions, including during colony development on agar surfaces and in high-density detergent-free shaken cultures. To address this hypothesis we used a previously described, hyper-aggregating suppressor strain of *M. smegmatis* that reports gene regulation responsive to cell-surface properties and/or biofilm microenvironments (14). The strain carries a suppressor mutation in the glycopeptidolipid biosynthesis gene (*mps*) in a Δ*lsr2* background that rescues the deficiencies in cell-cell adhesion and biofilm formation of the parent Δ*lsr2* strain(14). Importantly, the suppressor mutation does not rescue the expression of genes that are under negative regulation of Lsr2, implying that these genes are directly repressed by Lsr2 (14). Thus, the *lsr2*-independent genes responsive to the suppressor mutation are likely responding to signals sensing changes in environments that occur during aggregated growth such as pellicle development. We therefore reasoned that if *M. smegmatis* encodes regulators that recognize the *inlps* promoter and regulate its expression in a manner that responds to cellular growth in biofilms, then the regulation will be diminished in *lsr2* mutation, but restored back in the Δ*lsr2/* Δ*mps* suppressor. The expression pattern of a P*_inlps_-*Dendra2 reporter in *M. smegmatis* was similar to those observed in Mtb: induced expression was observed only during later stages (4-day or later) of pellicle development, and the induction was not apparent in planktonic cultures or colonies (Fig. 6A-C). Induced expression of Dendra2 reporter was not observed in impaired biofilms of Δ*lsr2*, but was substantially restored in the Δ*lsr2/* Δ*mps* suppressor (Fig. 6C). Together, the findings in *M. smegmatis* suggest that signals inducing *inlps* expression likely originate from the unique environment formed within biofilms. The P*_inlps_*-Dendra2 reporter therefore can serve as a marker for biofilm development.

**Figure 6:**
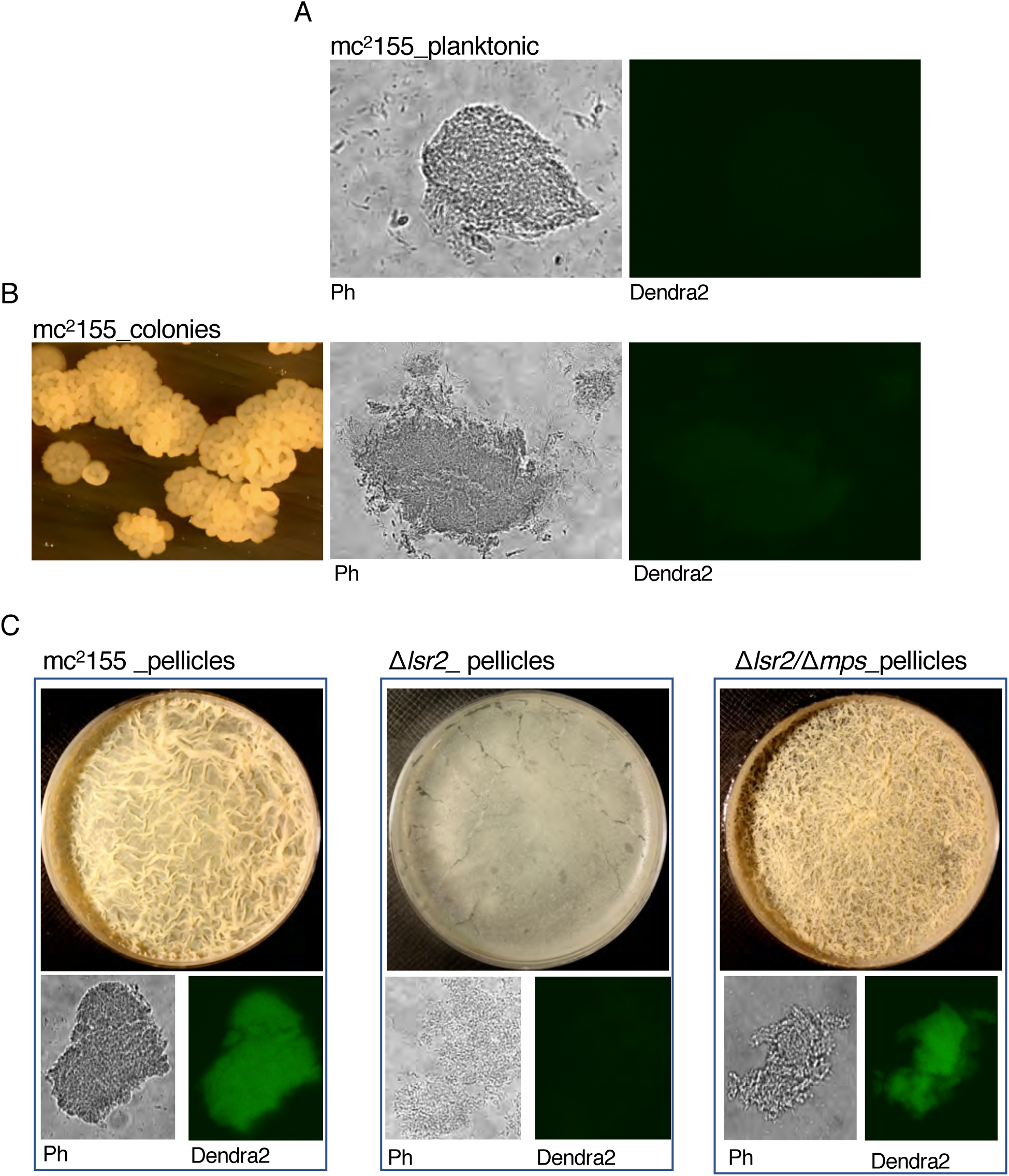
Expression pattern of INLPS promoter of Mtb in *M. smegmatis*. **A-C.** Reporter (P_inlps_**-**Dendra2) expression in recombinant *M. smegmatis* (mc^2^155) either in shaken detergent-free Sauton’s medium (planktonic) (A), or as colonies on Sauton’s medium agar plate (B, or as pellicle biofilms in static Sauton’s medium (C). Reporter strains of a biofilm-defective mutant Δ*lsr2* and its extragenic suppressor Δ*lsr2 /*Δ*mps* were also examined for P_inlps_**-**Dendra2 expression. Cells from each culture were smeared on a slide in 50% glycerol and imaged under wide-field fluorescent-light microscope at 20X magnification under phase contrast (Ph) or blue-filter (Dendra2). Morphologies of pellicles and colonies, from which cells were collected, are also shown. Cells in detergent-free shaken culture predominantly contain microscopic clumps (panel A).

## DISCUSSION

In this study we propose a model for elevated level of drug tolerance and high frequency of persisters of Mtb in biofilms. It is apparent from the Tn-seq analysis that significantly more genes are required for fitness of Mtb cells in biofilms than in other growth conditions. Cells growing within the complex biofilm architecture must be able to adapt to limiting nutrients and oxygen, as well as mechanical stresses. It is therefore not surprising that genes involved in the stringent response pathway (*relA*), cell wall integrity (*mma4*, *ponA2*, *mmpL10*, *fadA2*, *pks16 etc.)*, or nutrient homeostasis *(sugA*, *Rv2606c*, *sdaA*, *pstC2*, *pstA1*, *phoT*) are more crucial for Mtb fitness in biofilms than in planktonic culture. However, the correlation of a subset of these genes (*mma4* (31)*, ponA2* (41)*, pstA1* (40), Rv2224c (41)) with antibiotic tolerance in other studies suggests that a greater fraction of biofilm cells can survive for a longer exposure to antibiotics than planktonic cells. A logical outcome of the extended survival of the biofilm population can be hypothesized as an increase in the probability of occurrence of specialized persister cells, which emerge through mechanisms related to intrinsic defense against antibiotic-induced toxicity (42). In support of this hypothesis, we detected a lower frequency of persisters in morphologically normal biofilms of Δ*pstC2A1*, providing a reasonable platform to systematically evaluate the contribution of other identified genes in this study in drug tolerance of Mtb biofilms. We note that rifampicin sensitivity was also observed in the Δ*phoT and* Δ*dgt* mutants (JPR and AO, unpublished results). Moreover, the normal morphology of Δ*pstC2A1* biofilms suggests that the physical barrier presumably formed by the three-dimensional architecture of biofilms has limited contribution to the development of RIF tolerant persisters, instead the intrinsic sensitivity of resident cells to the drug play a significant role. Thus, a crucial role of the biofilm architecture is to create microenvironments that foster enrichment of drug tolerant cells.

A stringent genetic requirement for Mtb fitness in biofilms implies that conditions within biofilms are uniquely challenging. The unique environment of Mtb biofilms is clearly supported by the biofilm-specific induction of INLP. Further investigations on INLP function and its regulation will likely identify the conditions and signals that induce *inlps* expression within biofilms. SigC appears to be a strong candidate regulating the expression of INLPS in Mtb because it binds directly to the region upstream of Rv0096 (46), and overexpression of SigC induces the expression of Rv0096 (47). However, *M. smegmatis*, which also exhibits biofilm-specific upregulation of INLPS (Fig. 6), does not encode a SigC homolog, suggesting the likelihood of a functional equivalent of SigC in this species. Regardless, the fast-growing species offers a suitable model to identify the regulator and the signal inducing INLPS. INLP-like isonitrile compounds in *Streptomyces thioluteus* appear to be involved in copper transport (48), suggesting that INLP could be involved in metal transport in Mtb cells during biofilm development. Low intracellular zinc and nickel in a Δ*inlps* mutant of *M. marinum* (43) support INLP-dependent metal import in mycobacteria. Moreover, zinc is an important nutrient for Mtb biofilm formation (22). However, other functions of INLP cannot be ruled out, particularly because an intrinsic defect such as zinc import fails to explain why an *inlps* mutant could be trans-complemented by wild-type – as implied by the lack of *inlps* mutants in our Tn-seq screen.

Although direct evidence of Mtb biofilms during infection remain to be obtained, several genes required for fitness in biofilms are also important in Mtb survival during the early acute phase of infection –characterized by survival in macrophages (e.g. *ponA2*(41), Rv2224c(41, 49), *mma4*(50), *pstC2A1*(51)) – and in the chronic phase characterized by survival beyond 8 weeks of infection in a mouse model (e.g. *relA*(52)). This fits with an argument that *in vivo* biofilms of Mtb would likely represent a persistent community of bacilli that can successfully survive both innate and adaptive host resistance mechanisms. INLP has been implicated in Mtb survival during both acute and chronic phases of infection (53, 54), although it is unclear if the expression level of INLP varies during infection. It is tempting to speculate that a basal level of INLP expression has a different consequence on Mtb pathogenesis than an induced level. An INLP-based reporter could potentially address these possibilities and define *in vivo* biofilms of Mtb during infection.

## MATERIALS AND METHODS

### Bacterial strains and growth conditions

Unless otherwise indicated, an attenuated strain of Mtb, mc^2^7000(22), was used as the parent wild-type in the study. For planktonic cultures of mc^2^7000 or its recombinant strains, cells were grown at 37°C in Middlebrook 7H9 medium (Difco) supplemented with 10% (v/v) albumin dextrose catalase and oleate (OADC) (Difco), 0.05% (v/v) Tween-80 (Sigma), and 100 *µ*g/mL pantothenic acid (Sigma). For plate cultures, Middlebrook 7H11 agar supplemented with 10% OADC, and 100 *µ*g/mL pantothenic acid, or Sauton’s medium with 1% agarose and 100 *µ*g/mL pantothenic acid, were used. As necessary, zeocin, kanamycin, and hygromycin were added at concentrations of 25 *µ*g/mL, 20 *µ*g/mL, and 50*µ*g/ml, respectively while selecting the recombinant strains. For pellicle biofilms, logarithmic phase planktonic cultures of the tested strains were washed twice and then re-suspended in detergent-free 7H9 or Sauton’s media and diluted 1:100 into the corresponding detergent-free medium. Pellicles were grown at 37 °C with 4 mL per well in 12-well polystyrene tissue culture plates wrapped in parafilm for 5 weeks. For colonies grown for the Tn-seq, 1:10 dilutions of washed and re-suspended cultures were spotted onto 13mm Whatman polycarbonate membranes (Millipore Sigma, Cat. No. WHA110407), which were then dried for 1 hour to allow cells to attach. Inoculated membranes were placed on sterilized stacks of cardstock soaked in a pool of either detergent-free 7H9 medium or Sauton’s medium inside of a 100 x 35 mm polystyrene culture plate, and then grown at 37 °C for the indicated period of time. Medium was replenished as needed. For *Mycobacterium smegmatis*, mc^2^155 strain (wild-type) was grown at 37 °C in Middlebrook 7H9 medium supplemented with 10% (v/v) albumin dextrose and catalase (ADC) (Difco), and 0.05% (v/v) Tween-80, or in detergent-free Sauton’s medium. Agar plates of these medium were used to obtain colonies. For selection of *M. smegmatis*, 150 *µ*g/mL of hygromycin or 25 *µ*g/mL of zeocin was used as necessary. *M. smegmatis* pellicle biofilms were grown in Sauton’s medium, as previously described(14), by inoculating 10 μL from a saturated planktonic culture into 10 mL of detergent-free Sauton’s medium in 60mm polystyrene culture plates and incubating at 37 °C for 4 days. For molecular cloning, *Escherichia coli* GC5 cells were grown at 37 °C in Luria Broth or on LB agar under antibiotic selection conditions; 100 *µ*g/mL of carbenicillin, 50 *µ*g/mL of kanamycin, 150 *µ*g/mL of hygromycin or 25 *µ*g/mL of zeocin.

### Tn-seq screen and analysis

A transposon insertion mutant library of mc^2^7000 consisting of approximately 100,000 isolated mutant colonies was constructed using ΦMycoMarT7 bacteriophage carrying the Himar-1 transposon. The library was grown in 7H9OADC either as planktonic suspension to mid-logarithmic phase, in pellicle biofilms for 5-weeks, or as colonies on polycarbonate membrane for 18 days. Cells from each growth model, including planktonic suspension, were harvested and resuspended in PBS with 0.25% (v/v) Tween-80, vortexed for 30 seconds to disperse biomass, and sonicated in a Bath Sonicator (Branson) for 10 minutes. Ten-fold dilutions of bacteria from each sample were then plated out on ten 100mm 7H11OADC plates per replicate and incubated for 21 days. Colonies were harvested with a cell scraper and genomic DNA was extracted for further processing as described(34). Briefly, 2-5 *µ*g of genomic DNA was sheared with a Covaris M220 Focused-ultrasonicator (Covaris) into 400-600bp fragments, which were resolved by gel electrophoresis. The resolved fragments were gel-extracted (Qiagen Cat. No. 28606), and end-blunted using the Epicentre End Repair kit (Cat. No. ER0720). The fragments were then adenylated at the 3’ sequence end and custom adapter oligonucleotides ligated using the Epicentre FastLink DNA Ligation kit (Cat. No. LK6201H). Two-step, hemi-nested PCR-amplification of DNA fragments with a mixture of four staggered primers with homology to the end region of the transposon insertion resulted in a library of PCR amplicons at the junction site of transposon insertion and genomic DNA. Primers used for hemi-nested PCR and DNA sequencing are listed in Table S5. PCR with a set of short primers was performed with settings: 95 °C for 5 minutes; 20 cycles of 95 °C for 30 seconds, 58 °C for 30 seconds, and 72 °C for 45 seconds; 72 °C for 5 minutes. This was followed by hemi-nested PCR with a staggered primer set at 95 °C for 5 minutes; 10 cycles of 95 °C for 30 seconds, 58 °C for 30 seconds, and 72 °C for 45 seconds; 72 °C for 5 minutes. The library, prepared from a pool of three biologically independent sources of genomic DNA for every growth model, were sequenced at the transposon junction site using the Illumina Hi-Seq 2500 platform (TnSeq). Two biologically independent sets of libraries were sequenced for each growth model. The sequencing results were analyzed by TRANSIT(35) pipeline, using Terminal Total Reads (TTR) as the normalization method, to identify ORFs with significant underrepresentation in either pellicle or colony biofilms, relative to planktonic cultures.

### Plasmids and mutant constructions

Strains and plasmids used in this study are listed in Tables S6 and S7, respectively. A modified recombineering method, as described previously(14), was used for constructing isogenic deletions in the indicated genes. For allelic exchange substrates (AES) of a target gene, 250bp PCR amplicons corresponding to upstream and downstream of the gene were joined to the either side of a loxP-flanked zeocin-resistance cassette by sewing-PCR. Oligonucleotides used for AES generation are listed in Table S5. The AES were electroporated into a recombineering strain of mc^2^7000 carrying the plasmid pJV53-SacB (14), and plated on 7H11 agar with 25 μg/mL of zeocin and 20 *µ*g/mL kanamycin. The genotype of zeo^r^ colonies was confirmed by PCR. The recombineering plasmid, pJV53-SacB, was removed by selecting the mutant strains on 7H11 agar with 15% sucrose and 25 *µ*g/mL zeocin. The sucrose-resistant colonies were screened for kanamycin sensitivity. A second PCR was performed to confirm gene deletion gene using primers homologous to upstream and downstream ends of AES and to the zeocin-resistance cassette. As necessary, mutants were complemented by the corresponding gene cloned along with a 500bp upstream promoter region, or fused to *hsp60* promoter, on an integrative plasmid (pMH94). For a reporter strain of a mutant, an integrative plasmid expressing mCherry from the constitutive *hsp60* promoter was electroporated into the mutant and transformants were selected on 7H11OADC plates with 25 *µ*g/mL zeocin and 20 *µ*g/mL kanamycin. For construction of INLP expression reporter, the putative promoter and regulator elements of INLP harbored in the 500bp upstream of the gene cluster were amplified using the primers listed in Table S5 and cloned upstream of Dendra2 at XbaI and NdeI restriction sites in the plasmid pYL026. The resulting plasmid, pJR36, was electroporated into mc^2^155, mc^2^7000 and Mtb (Erdman) strains for analysis of the reporter expression.

### Microscopic imaging and analysis

For imaging on a widefield fluorescent microscope, Nikon Eclipse TE2000-E microscope with either a 20x (NA: 0.75) objective was used. Images were taken with a 600nm exposure using Pro-Image 7.0 software under transmitted light (phase contrast), or with X-Cite 120 Fluorescence Illumination System with a blue GFP(R)-BP filter (Excitation: 460-500nm; BA: 510-560nm) for Dendra2 fluorescence or a yellow Y-2E filter (Excitation: 540-580nm; BA:600-660nm) for mcherry. CSLM imaging was done with a Leica SP5 confocal microscope with a 488nm laser for Dendra2 and a 593nm laser for mcherry, under the settings described. All post-acquisition analysis was done using FIJI image analysis software. All settings on the microscopes as well as in FIJI were maintained across compared samples during image acquisition and analysis.

### Analysis of persisters in pellicle biofilms

Biofilms of the indicated strains were grown in pellicles as described above, and then exposed to 50 *µ*g/mL of RIF (or an equal volume of DMSO as a control) for 7 days. After exposure, pellicles were collected and washed 3 times with 1xPBS with 0.25% tween-80 to remove residual antibiotic, and then left on a shaker at 4°C overnight in the presence of sterile 10mm glass beads. Serial dilutions were plated on 7H11 agar to enumerate the viable colonies.

### Gene expression analysis by RNA-seq and RT-PCR

Using a Qiagen RNeasy kit (Cat. No. 74104), total RNA from desired cultures of the indicated strains of Mtb were isolated from 20 mL cultures obtained from mid-exponential growth phase in 7H9 medium. For biofilm transcriptomics, mc^2^7000 was grown planktonically to mid-exponential phase in detergent-free Sauton’s medium or into pellicles for 5-weeks in detergent-free Sautons’s medium as described above. The pellicle biomass portion was scooped out of the liquid medium using parafilm. RNA was extracted from the pellicles using RNAEasy kit (Qiagen Inc.). Contaminating genomic DNA from the RNA preparations was removed with the Thermo Fisher Scientific Turbo DNA-free Kit (Cat. No. AM1907). The ribosomal RNA from a total of 5 μg RNA was removed with the Illumina RiboZero Kit (now discontinued). Strand-specific cDNA libraries were prepared from 100ng mRNA of each sample using the Illumina Scriptseq v2 Complete Kit (Cat. No. SSV21106). Libraries were sequenced on the Illumina NextSeq500 platform at the Wadsworth Center, and results were analyzed by Rockhopper(55) at the default settings using the MTB H37Rv reference genome (NC_000962). All oligonucleotides used for RT-qPCR are listed in Supplementary Table S5. For RT-qPCR, DNA-free RNA was extracted from desired bacterial cultures as described above for RNA-seq. Using the Fisher Maxima First Strand cDNA Synthesis Kit (Cat. No. K1641), cDNA was generated from 200 ng of total RNA from each specified sample. RT-qPCR reactions were prepared using 2 μL of cDNA reaction mixture, 0.5 μL from 10 μM stock for each gene-specific primer per reaction, and Applied Biosystems SYBR Green Master Mix (Cat. No. A25742) as per the manufacturer’s instruction. qPCR was performed on an Applied Biosystems 7500 Fast Real-Time PCR System (Cat. No. 4351106) using the cycle conditions: 95°C for 10 minutes; followed by 40 cycles of 95°C for 20 seconds, and then 60°C for 1 minute.

Primers corresponding to SigA transcripts were used as an endogenous control. Reactions without the cDNA were used as No-template negative control. Expression of a target gene (tgene) in pellicles relative to planktonic was calculated as 2^-^[^ΔCt(pellicle_tgene – pellicle_sigA)-ΔCt(plnk_tgene – plnk_sigA)^].

### INLP analysis

Pellicle biofilms or planktonic cultures were grown in detergent-free Sauton’s medium prior to crude lipid extraction. 1 mM of 2-^13^C-Gly (Sigma) was included in selective biofilm cultures for isotope incorporation. Cells from planktonic cultures were transferred to glass tubes and an equal volume of chloroform : methanol (2:1) was added to each culture, followed by intermittent vortexing for 30 seconds every 15 minutes for 2 hours. The mixture was then centrifuged at ∼900g for 10 minutes on a Thermo Scientific Sorvall Legend XTR (Cat. No. 75004521) centrifuge. The lower, organic phase was separated by Pasteur pipette and dried under N_2_ gas. Pellicle biomass was separated from the liquid portion of the cultures using parafilm. The biomass was washed and resuspended in 5 mL PBS. Equal volume of chloroform : methanol (2:1) was added to the suspension and the lipid in organic phase was collected and dried as described above. Dried crude extracts were dissolved in methanol to final concentration of 0.1mg/ ml. Samples were centrifuged at 12,000 rpm for 5 min, and the supernatant was used for LC-MS analysis.

#### Liquid chromatography-mass spectrometry (Orbi-trap)

Samples of extracted metabolites were analyzed using a liquid chromatography system (LC; 1200 series, Agilent, Santa Clara, CA) that was connected in-line with an LTQ-Orbitrap-XL mass spectrometer equipped with an electrospray ionization source (Thermo Fisher Scientific, Waltham, MA). The LC was equipped with a reversed-phase analytical column (length: 150 mm, inner diameter: 1.0 mm, particle size: 5 *µ*m, Viva C18, Restek, Bellefonte, PA). Acetonitrile, formic acid (Optima grade, 99.5+%, Fisher, Pittsburgh, PA), and water purified to a resistivity of 18.2 MΩ·cm (at 25 °C) using a Milli-Q Gradient ultrapure water purification system (Millipore, Billerica, MA) were used to prepare mobile phase solvents. Solvent A was 99.9% water/0.1% formic acid and solvent B was 99.9% acetonitrile/0.1% formic acid (v/v). The elution program consisted of isocratic conditions at 5% B for 2 min, a linear gradient to 98% B over 25 min, isocratic conditions at 98% B for 10 min, at a flow rate of 150 *µ*L/min. Full-scan, high-resolution mass spectra were acquired over the range of mass-to-charge ratio (*m*/*z*) = 70 to 1000 using the Orbitrap mass analyzer, in the positive ion mode and profile format, with a mass resolution setting of 100,000 (measured at full width at half-maximum peak height, FWHM, at *m*/*z* = 400). For tandem mass spectrometry (MS/MS or MS^2^) analysis, precursor ions were fragmented using higher energy collisional dissociation (HCD) under the following conditions: MS/MS spectra acquired using the Orbitrap mass analyzer, in centroid format, with a mass resolution setting of 7500 (at *m*/*z* = 400, FWHM), isolation width: 5 *m*/*z* units, normalized collision energy: 38%, default charge state: 1+, activation time: 30 ms, and first *m*/*z* value: 100. Mass spectrometry data acquisition and analysis were performed using Xcalibur software (version 2.0.7, Thermo). Comparative metabolomics analysis was performed using MS-DIAL(56).

#### Liquid chromatography-mass spectrometry (QTOF)

LC-MS analysis was performed using an Agilent Technologies 6520 Accurate-Mass Q-TOF LC-MS instrument and an Agilent Eclipse Plus C18 column (4.6 x 100 mm). A linear gradient of 2-98% acetonitrile (v/v) over 30 min in H_2_O with 0.1% formic acid (v/v) at a flow rate of 0.5 mL/min was used. HRMS/MS analysis was conducted using targeted MS/MS with collision energy of 20V.

#### Click chemistry analysis

Compound **1** and **2** were partially purified from the wild-type culture extract using HPLC. This was conducted using an Agilent 1200 HPLC with a Waters Atlantis T3 OBD column (10 × 250 mm) using a linear gradient of 5-95% CH_3_CN (v/v) over 30 min in H_2_O without formic acid at a flow rate of 3.5 mL/min. Fractions were screened using LC-MS as described above. Fractions containing **1** and **2** were combined, dried, and re-dissolved in methanol, followed by addition of 100 *µ*M of 3,6-Di-2-pyridyl-1,2,4,5-tetrazine and incubated at r.t. for 4.5 hours before LC-MS analysis.

## Acknowledgments

This work was supported by grants to AKO (NIH: AI132422, NIH: AI144474) and WZ (NIH: DP2AT009148, the Alfred P. Sloan Foundation, and the Chan Zuckerberg Biohub Investigator Program). Support from the Wadsworth Center core facilities– Fluorescent and Light Microscopy Imaging and Analysis, and Applied Genomics Technology– are acknowledged. Authors acknowledge Yong Yang and Jennifer Gundrum for technical help. Authors are also thankful to Christopher Sassetti and Richard Baker for help in Tn-seq experiments, William Jacobs Jr. for the gift of *inlps* cosmid, and Anthony T. Iavarone for assistance with LC-MS (Orbi-trap) analysis. Support to The QB3/Chemistry Mass Spectrometry Facility, University of California, Berkeley by NIH grant (1S10OD020062) is acknowledged.

## Authors Contribution

JPR and AKO designed and performed experiments, as well as analyzed data on all experiments except chemical analysis of INLP. WC, NAZ and WZ designed and performed experiments, and analyzed data on INLP analysis. JPR, WC, NAZ and WZ wrote the manuscript.

## Financial Conflict of Interest

Authors declare no financial conflict of interest.

## Supplemental Information

**Figure S1:**
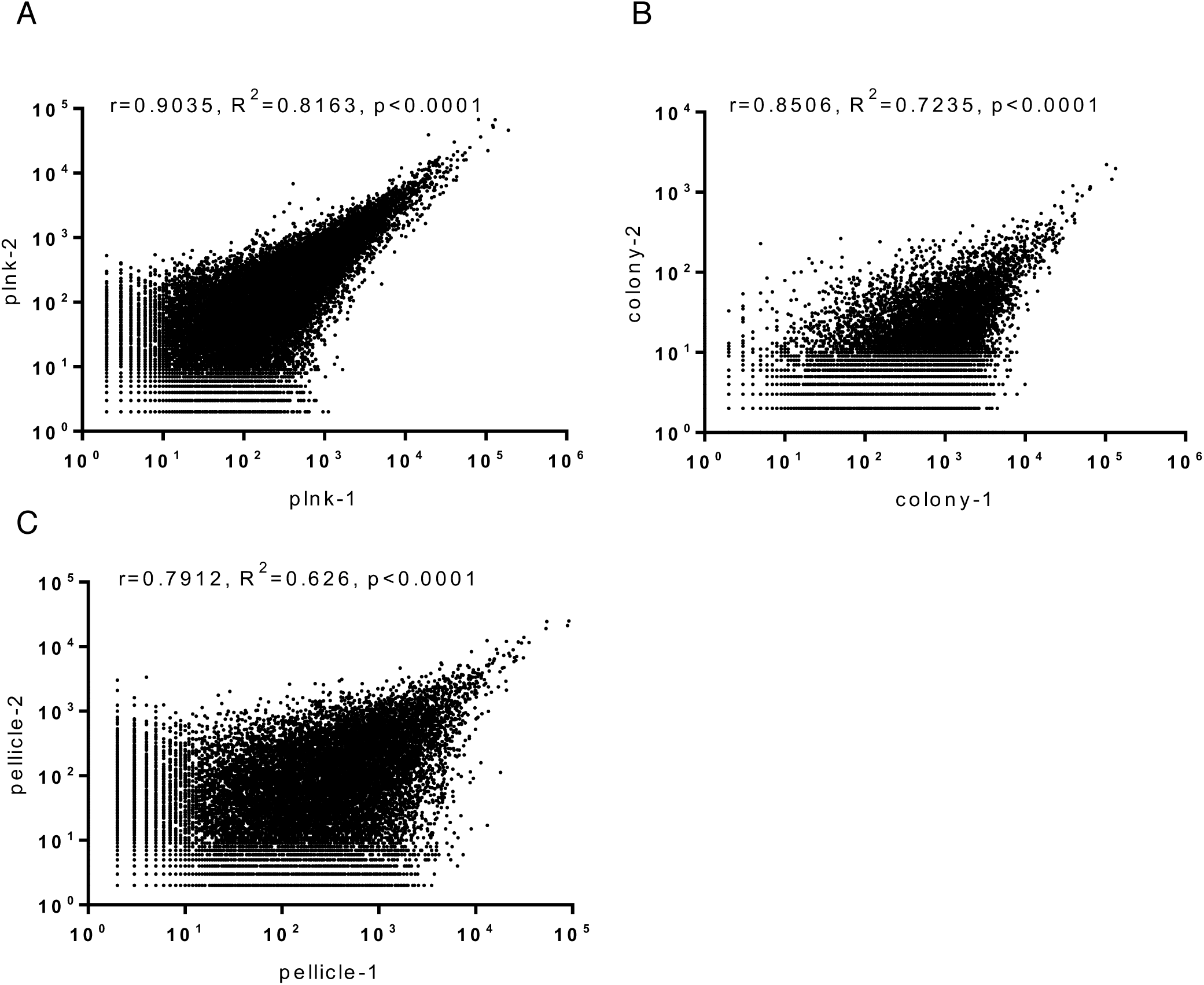
Tn-Seq of Mtb biofilms. **A-C.** Correlation between two Tn-Seq replicates of planktonic (plnk) (A), colonies (B), and pellicle biofilms (C). Number corresponding to sequencing reads mapped to each transposon insertion site in a sample is plotted against its replicate. Potential insertion sites (TA dinucleotides) that contained zero or only one mapped read in both replicates were excluded from the correlation analysis. Any remaining insertion sites that had zero mapped reads in one of the replicates was converted to one, to allow for log-scale plotting. The Pearson coefficient (r), R-squared values, and statistical significance of correlation were calculated in GraphPad Prism 7. See Table S1 for the summary of metadata.

**Figure S2:**
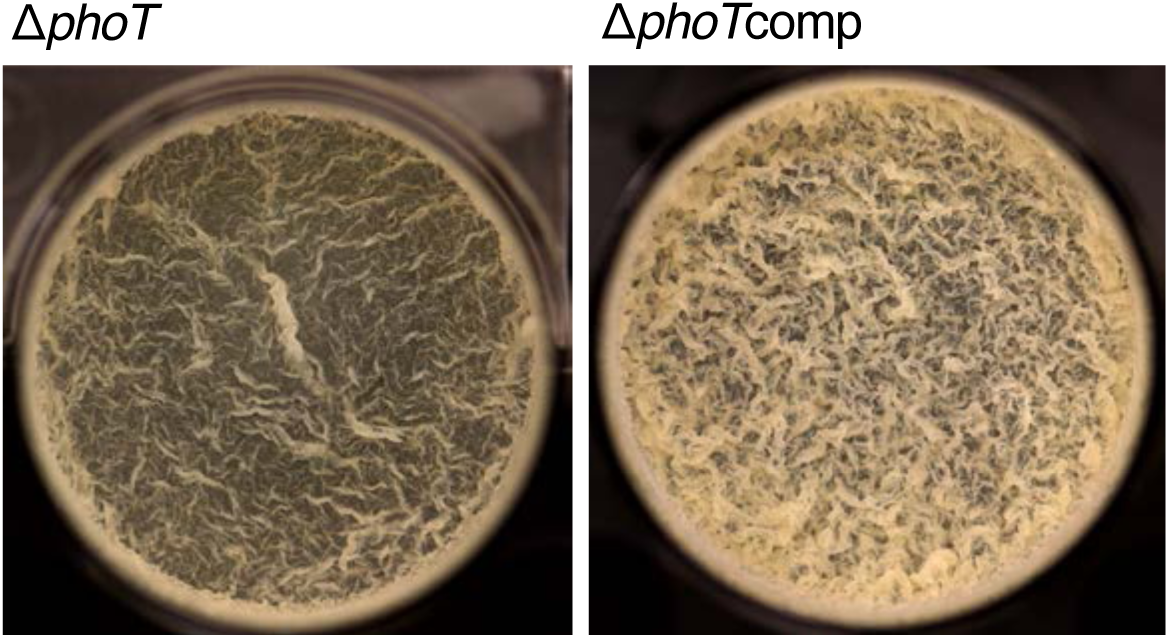
Pellicle development in Δ*phoT* upon extended incubation. A top-down view of pellicles of Δ*phoT* and its complemented strain at the liquid medium interface after six weeks of incubation.

**Figure S3:**
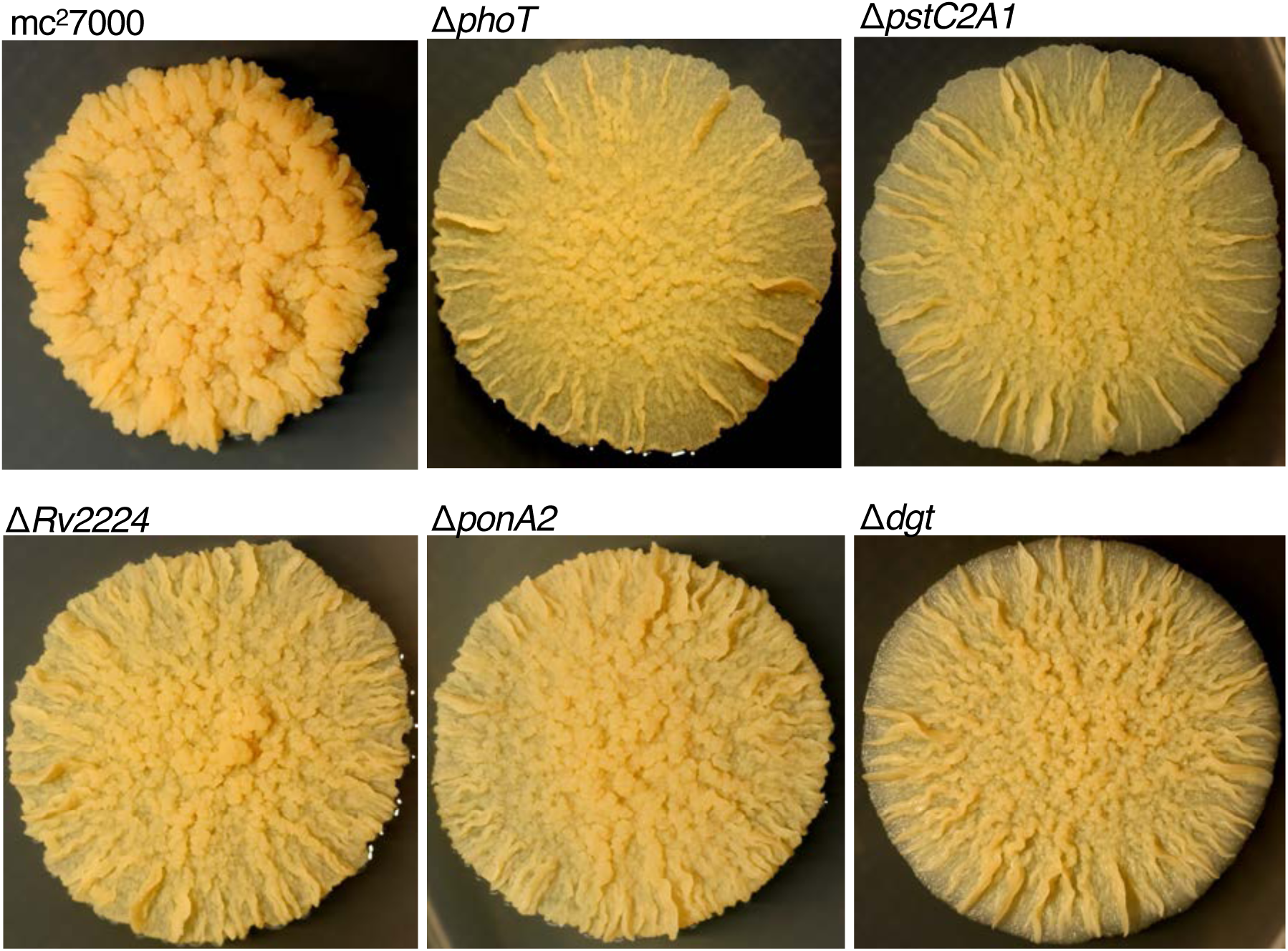
Morphologies of colonies formed by the indicated mutants after six weeks of incubation on Sauton’s medium based agar plates.

**Figure S4:**
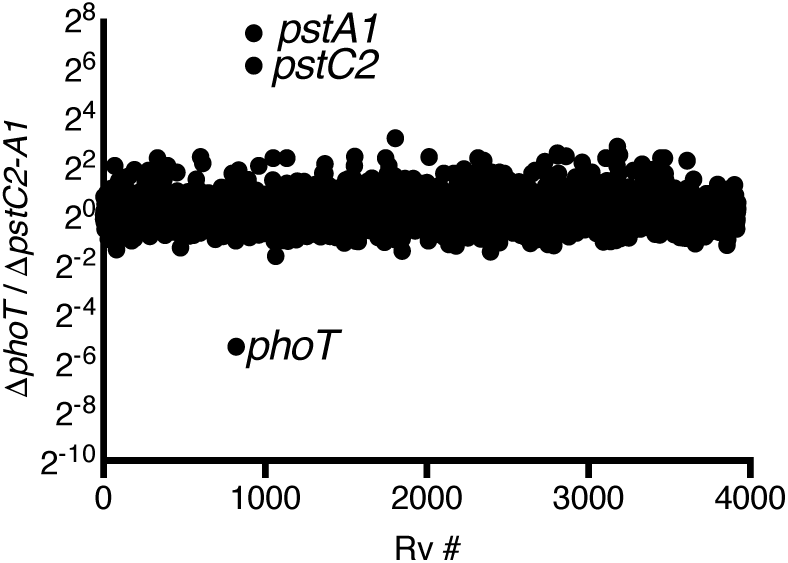
Similarity between transcriptomes of Δ*phoT and* Δ*pstC2A1* mutants. Ratio of the reads corresponding to each Rv # between Δ*phoT and* Δ*pstC2A1* indicate a broad similarity in transcriptomes of the two mutants, while each being similarly different from the wild type. The only significant differences (> 4-fold) are in the genes deleted in the corresponding mutants. The RNA-seq data are provided in Table S3.

**Figure S5:**
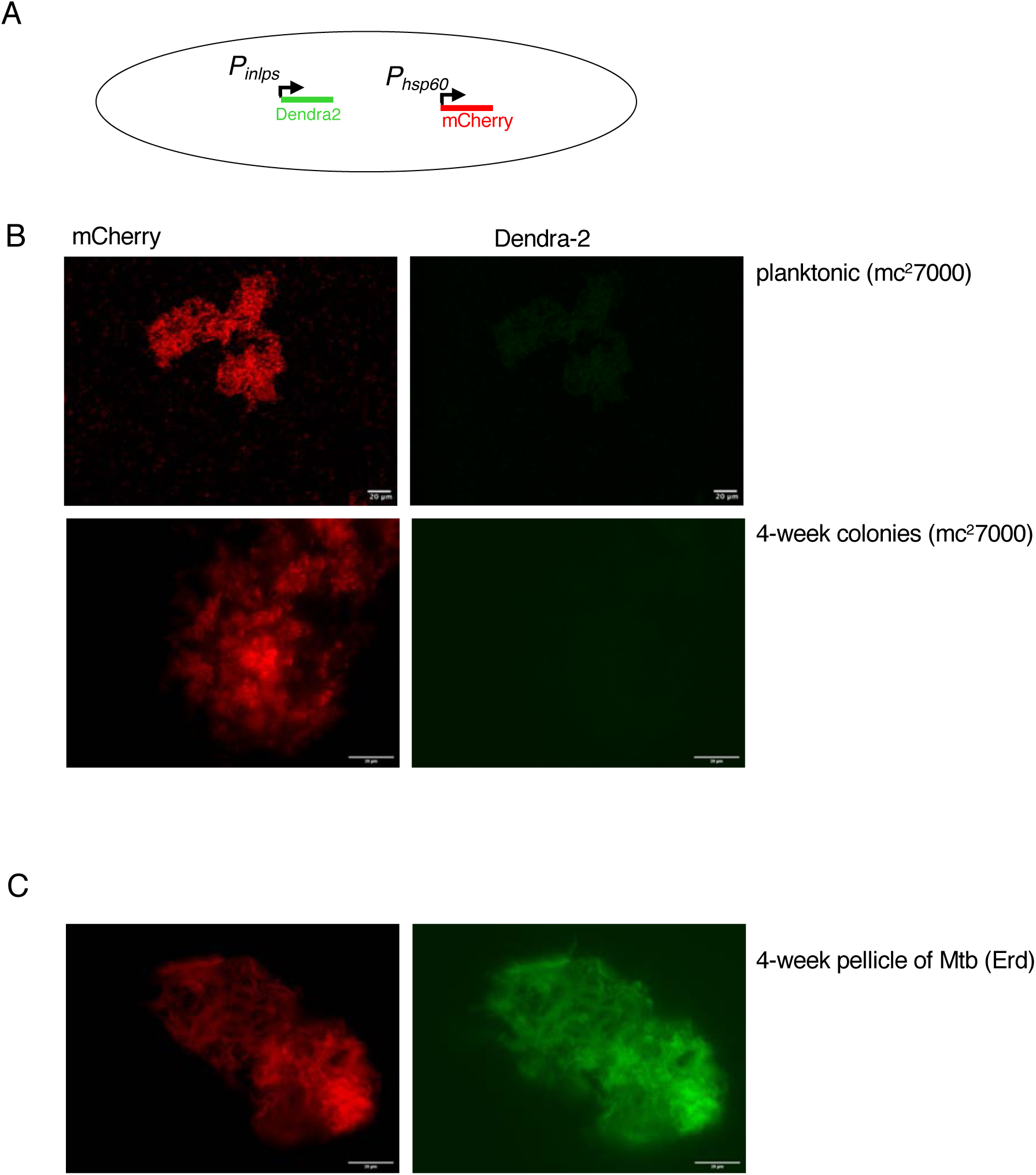
Induced expression of INLPS in Mtb biofilms. **A.** A schematic of the reporter construction for strains used in panels b, c, and in Figure 4c of the paper. **B.** Lack of INLPS expression in detergent-free planktonic mc^2^7000 cells after 2-weeks of incubation, and in 4-week colonies. **C.** Expression of INLPS in 4-week pellicles of Mtb (Erdman). Clump of cells of the indicated strains from either colonies or pellicles were collected, fixed in formalin, smeared on slides in 80% glycerol and visualized under either confocal (panel b planktonic) or widefield fluorescence microscope at 20X magnification. Scale bar represents 20 *µ*. Images were analyzed by FIJI. Confocal images are shown as maximum projection across stacks.

**Figure S6:**
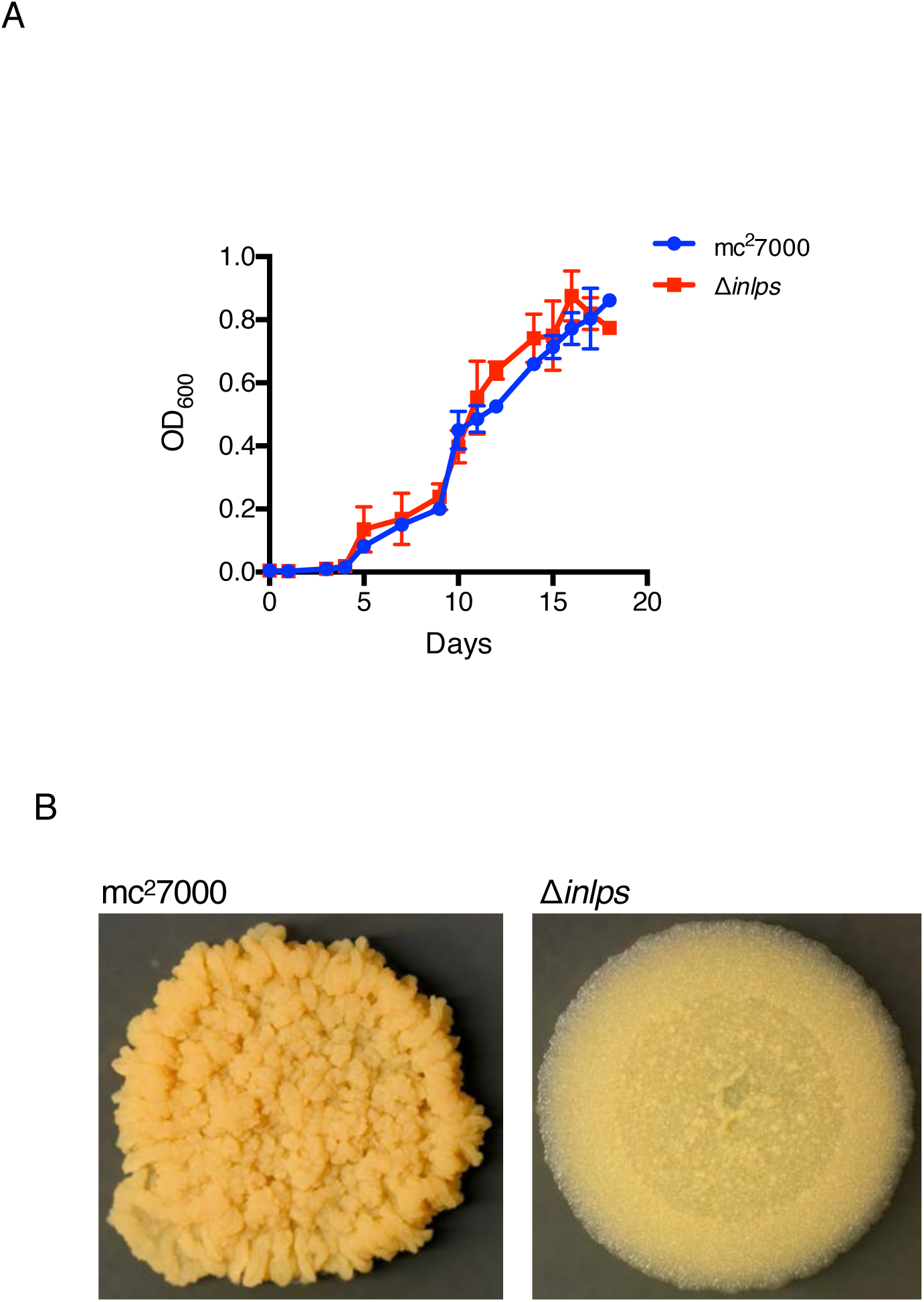
Growth of mc^2^7000 and its isogenic Δ*inlps* in planktonic culture (A), and on agar plates in Sauton’s medium (B).

**Figure S7:**
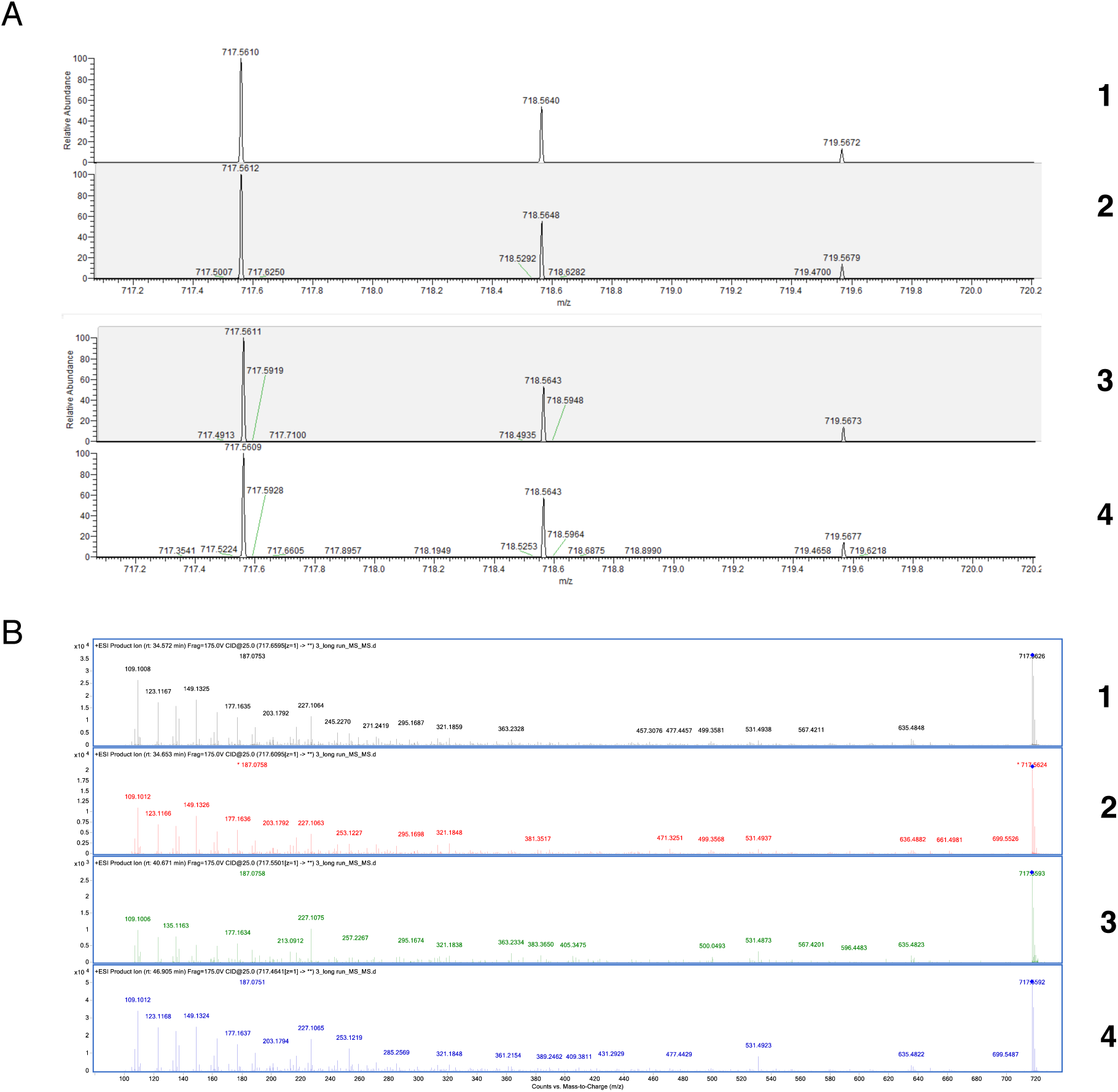
High-resolution mass spectrometric (HRMS) analysis of INLP isomers**. A.** HRMS analysis of metabolites **1-4** shown in figure 5 (proposed molecular formulas C_40_H_73_N_6_O_5_^+^, Δ = −3.4 ppm). **B.** MS/MS fragmentation spectra of **1**-**4** showing nearly identical patterns. The presence of lipid chains are also suggested from these MS/MS spectra.

**Figure S8:**
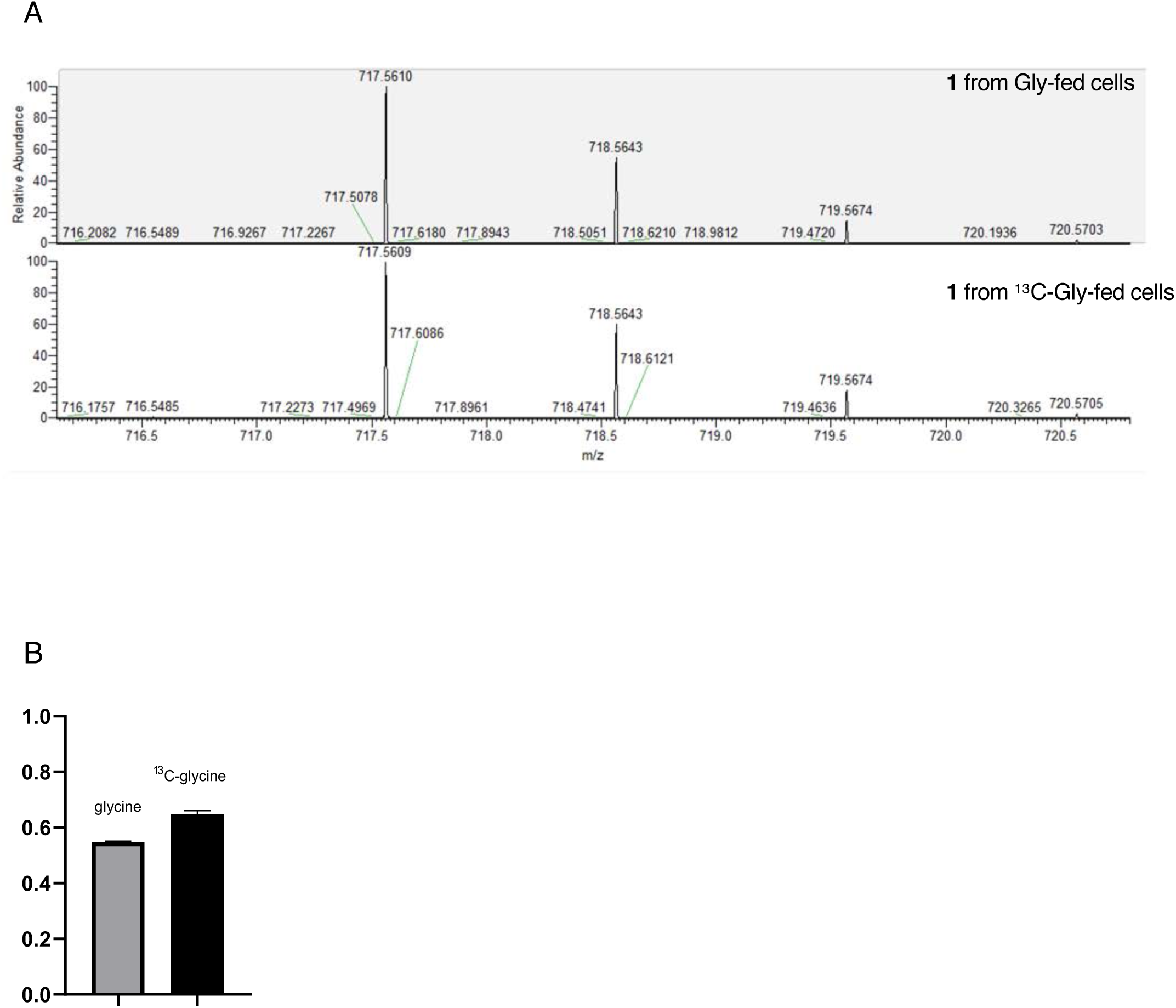
HRMS analysis of INLP isomer **1** upon Gly feeding. **A**. Feeding of 2-^13^C-Gly to the culture of Mtb demonstrated that C(2) of Gly was weakly but consistently (∼10%) incorporated into **1** during biofilm formation. **B.** Isotopic incorporation was calculated using the ratios of integrated area under curve (AUC) of [M+2] over [M+1]. The ratios were 55% and 65% for Gly and 2-^13^C-Gly feeding samples, respectively. Similar incorporation was observed for INLP isomers **2**-**4**.

## Attached data files

**Table S2:** Tn-seq based assessment of the impact of each gene on growth of *M. tuberculosis* (mc^2^7000) in pellicle- and colony-biofilms, relative to planktonic culture.

**Table S3:** RNA-seq based comparison between transcriptomes of mc^2^7000, Δ*phoT* and Δ*pstC2-A1*.

**Table S4:** RNA-seq based comparison between planktonic and biofilm transcriptomes of mc^2^7000.

**Table S5:** List of oligonucleotides used in this study.

**Table S6:** List of bacterial strains used in this study.

**Table S7:** List of plasmids used in this study.

